# A novel variant in *ADPRS* disrupts ARH3 stability and subcellular localization in children with neurodegeneration and respiratory failure

**DOI:** 10.1101/2024.06.14.597428

**Authors:** Maxwell Bannister, Sarah Bray, Anjali Aggarwal, Charles Billington, Hai Dang Nguyen

## Abstract

**Purpose:** ADP-ribosylation is a post-translational modification involving the transfer of one or more ADP-ribose units from NAD+ to target proteins. Dysregulation of ADP-ribosylation is implicated in neurodegenerative diseases. Here we report a novel homozygous variant in the *ADPRS* gene (c.545A>G, p.His182Arg) encoding the mono(ADP-ribosyl) hydrolase ARH3 found in 2 patients with childhood-onset neurodegeneration with stress-induced ataxia and seizures (CONDSIAS).

**Methods:** Genetic testing via exome sequencing was used to identify the underlying disease cause in two siblings with developmental delay, seizures, progressive muscle weakness, and respiratory failure following an episodic course. Studies in a cell culture model uncover biochemical and cellular consequences of the identified genetic change.

**Results:** The ARH3^H182R^ variant affects a highly conserved residue in the active site of ARH3, leading to protein instability, degradation, and reduced expression. ARH3^H182R^ additionally fails to localize to the nucleus. The combination of reduced expression and mislocalization of ARH3^H182R^ resulted in accumulation of mono-ADP ribosylated species in cells.

**Conclusions:** The children’s clinical course combined with the biochemical characterization of their genetic variant develops our understanding of the pathogenic mechanisms driving CONDSIAS and highlights a critical role for ARH3-regulated ADP ribosylation in nervous system integrity.

## INTRODUCTION

ADP-ribosylation is the post-translational addition of ADP-ribose (ADPr) to protein targets identified as early as 1963.^1^ The following decades unraveled its pervasive roles in integral cellular processes such as transcription, DNA replication, protein turnover, DNA damage repair, immune responses, and apoptosis.^2^ This diversity in function is enabled by a family of 17 different ADP-ribosyl transferases (ARTs). ARTs can catalyze the addition of a mono-ADP-ribose unit (MAR and MARylation) on target substrates that can subsequently be extended to form long, branching chains of poly-ADP-ribose (PAR and PARylation). The lengths and linkage patterns of ADPr units on protein substrates can alter interaction affinities, localization, and activity.^3^ Additionally, ARTs can modify a chemically diverse set of amino acid residues, including serine, aspartate, lysine, arginine, and cysteine.^4–6^ Recent work extended the functional diversity of ADP-ribosylation showing not only that DNA and RNA can be modified with ADPr, but also that free ADPr polymers are signaling molecules in parthanatos, an ADPr-mediated form of cell death.^7–9^

These critical functions of ADP-ribosylation necessitate tight regulation. The principal ADP-ribosylhydrolase in human cells is poly(ADP-ribosyl) glycohydrolase (PARG) and can rapidly remove PAR.^10^ However, PARG is unable to cleave the linkage between target proteins and the proximal ADPr unit.^10^ Instead, the final ADPr can be removed by various other mono-ADP-ribosylhydrolases with different specificities for the modified amino acid residue.^11,12^ ADP-ribosylhydrolase 3 (ARH3) is the only known hydrolase that can remove serine-MAR.^10^ ARH3 can remove PAR in an exo-manner and entire chains in an endo-manner like PARG.^11^ However, the vastly greater processivity of PARG means that the observed effect of ARH3 on PAR levels is largely from its removal of serine-MAR. Serine-MARylation is the rate-limiting step in a form of ADP-ribosylation signaling catalyzed by the PARP1:HPF1 complex recently shown to dominate in response to DNA damage.^13^ Finally, ARH3 is known to degrade protein-free PAR chains that are cleaved in an endo-manner by PARG^14^, and the accumulation of these PAR chains in the cytoplasm is a known step of parthanatos.^7^

Variants in the gene *ADPRS* (also known as *ADPRHL2*, HGNC:21304) encoding ARH3 were first reported in 2018 as causing an autosomal recessive neurodegenerative disease presenting in previously healthy children as progressive ataxia, seizures, loss of developmental milestones, cerebellar atrophy, and peripheral neuropathy.^15,16^ The most severe symptoms (seizures, death) occur in an episodic manner often preceded by some physiological stressor, most commonly febrile illness but as diverse as a near-drowning event.^15^ This disorder has been termed Childhood-Onset NeuroDegeneration, Stress-Induced, with variable Ataxia and Seizures (CONDSIAS) (MIM #618170). Emerging from the identification of approximately 40 different variants in CONDSIAS is a story of loss of ARH3 function followed by steady-state accumulation of Ser-MAR and, to an even greater extent, following cellular damage.^14–26^

In this study, we identified a novel homozygous *ADPRS* variant (c.545A>G, p.His182Arg, H182R) in two siblings with a clinical picture consistent with CONDSIAS. Using *in vitro* cell models, we found that the H182R variant conferred loss of ARH3 function. ARH3^H182R^ is rapidly degraded in cells, resulting in a failure to resolve accumulated MAR in ARH3^KO^ cells. Moreover, the residual ARH3^H182R^ in cells localized to the cytoplasm but not the nucleus, further preventing suppression of nuclear MARylated substrates.

## MATERIALS AND METHODS

### Genome sequencing

Clinical genetic testing was performed according to protocols standard in the University of Minnesota Molecular Diagnostic Laboratory. Briefly, genomic DNA was extracted using the QIAamp DNA Blood Midi Kit or QIAamp DNeasy Blood & Tissue Kit (QIAgen). Genome sequencing libraries were made using Illumina genome DNA prep reagents (Illumina) and sequenced on an Illumina Novaseq instrument with paired-end 150 base-pair reads. Reads were mapped to GRCh37 using the BWA algorithm, and variant calling was performed with the Genome Analysis Toolkit (GATK) version 4.1 genotyper.

### Confirmatory Sanger sequencing

Sanger sequencing was performed using standard approaches. Primers were designed to amplify and sequence *ADPRS* NM_017825.3: c.545A>G (p.His182Arg). Following amplification of the genomic region containing this specific variant from genomic DNA, bi-directional Sanger sequencing was performed. Analysis of family members for *ADPRS* c.545A>G included use of a positive, affected proband as a positive control.

### Antibodies, oligos, and plasmids

All antibody information is available in **Table S1**. All primers and plasmids used in this study are available in **Table S2** and **S3**, respectively.

### Plasmids

The ARH3^WT^-Flag and ARH3^H182R^-Flag were synthesized by Gene Universal and subcloned into pDonor221 to generate pEntry plasmids prior to Gateway cloning into the appropriate pDest plasmids (pLenti6.2-DEST, Addgene #87071, for lentivirus, pDest17 for protein expression). These ARH3-Flag coding sequences carry a stop codon upstream from the 3xFLAG-V5 tags within the pLenti6.2 plasmid. All plasmids were confirmed by Sanger sequencing. Guide-RNA targeting the *ADPRS* gene were cloned into PX459 plasmid (Addgene #62988).

### Cell culture and transfection

U2OS and its derivative cell lines were maintained in DMEM (Corning #MT15013CM) supplemented with 10% FBS (Corning, #MT35011CV), penicillin-streptomycin (100 U/mL, Gibco #15140122), and L-glutamine (2 mM, Gibco #25030081). To knockout the *ADPRS* gene, U2OS cells were transfected with the PX459 plasmid containing sgARH3-2 (**Table S2**) using the Lipofectamine 3000 transfection reagent (Invitrogen #L3000015) according to the manufacturer’s instructions. Transfected cells were selected in puromycin (1 μg/mL, Gibco #A1113803) for 3 days. Knockout efficiency was determined by western blot.

### Lentivirus production and transduction

For lentivirus production, HEK293T cells (Clontech) were co-transfected with the packaging plasmids pCMV-dR8.2 dVPR (10 μg, Addgene #8455) and pCMV-VSV-G (10 μg, Addgene #8454) along with a pLenti6.2 expression plasmid (10 μg, Addgene #87071) carrying ARH3^WT^-Flag or ARH3^H182R^Flag using the calcium phosphate-mediated ProFection Mammalian Transfection System (Promega #E1200) for 72 hours. HEK293T cell confluency was maintained at 60% of a 10 cm plate at the time of transfection, and media was changed 7 hours post-transfection. Supernatants containing virions were collected at 72 hours post transfection. Pooled U2OS *ADPRS*-knockout cells were transduced using the spinoculation method in the presence of polybrene (4 μg/mL, Millipore #TR-1003-G) and then cultured in 1 μg/mL blasticidin (VWR #97064-358) for 5 days. Clonal lines were expanded by limiting dilution and confirmed to express Flag-tagged ARH3 by immunoblotting.

### Western blot

Cells were lysed in a lysis buffer (100 mM Tris pH 6.8, 1% SDS) solution and denatured at 95°C for 5 min. Protein concentrations were quantified using the Pierce BCA Protein Assay Kit (Thermo Scientific #23227) and mixed 1:1 with 2× SDS-PAGE loading buffer (100 mM Tris at pH 6.8, 12% glycerol, 3.5% SDS, 0.2 M DTT). Samples were run on polyacrylamide gels, transferred onto PVDF membranes, and then blocked in Tris-buffered saline containing 0.05% Tween-20 (TBS-T) and 5% milk for 1 h at room temperature. Membranes were then immunoblotted with primary antibodies (**Table S1**) overnight at 4°C. Membranes were washed 3 times with TBS-T buffer and incubated for 1 h at room temperature with secondary antibodies conjugated to horseradish peroxidase. Membranes were washed 3 times for 10 min with TBS-T and developed with an enhanced chemiluminescence (ECL Bio-Rad 1705061) substrate. Signals were detected using the ChemiDoc imaging system (Bio-Rad) and analyzed using BioRad ImageLab v6.0.1. software.

### Immunofluorescence

For immunofluorescent staining of MAR, U2OS cells were extracted 1x PBS containing 0.2% Triton-X100 supplemented with PARP inhibitor (olaparib, 1 μM, Selleckchem #S1060) and PARG inhibitor (PDD 00017273, 1 μM, Tocris #5952) to stabilize ADP-ribosylated species for 5 min on ice prior to fixation with 3% paraformaldehyde/2% sucrose for 15 min at ambient temperature. Subsequently, cells were permeabilized with PBS containing 0.2% Triton-X100 for 10 min, blocked in blocking buffer (1x TBS containing 5% BSA, 0.05% Tween-20) for 1 hr, and incubated in MAR primary antibody (1:500, Bio-Rad #AbD33204) in blocking buffer overnight at 4°C. Next day, cells were washed 3 times with PBS-T before incubation with Alexa Fluor 488 anti-rabbit secondary antibody for 1 hr at ambient temperature and DAPI staining.

For cell-cycle analysis of U2OS cell lines, a quantitative image-based cytometry (QIBC) method was used. Briefly, cells were labeled with 10 μM EdU for 30 min and processed with the Click-IT EdU Alexa Fluor 488 Imaging Kit (Invitrogen, #C10337) according to the manufacturer’s instructions.

For immunofluorescence analysis of ARH3, U2OS cells were fixed with 3% paraformaldehyde/2% sucrose for 15 min at ambient temperature, permeabilized with 100% methanol on ice for 10 min, blocked in 2% BSA in PBS, and incubated for 12 hr at 4°C in the presence of mouse anti-FLAG (1:1000, Sigma #F1804) and rabbit anti-CD40 (1:500, Abcam #ab224639). Cells were then washed 3 times with 0.2% BSA in PBS before incubation with Cy3 anti-rabbit and Cy5 anti-mouse secondary antibodies for 1 hr at ambient temperature. Images were captured using a Leica DMi8 microscope.

### Immunofluorescence image analysis

Image segmentation of nuclei and whole cells was performed using the cellpose algorithm implemented in Python.^27^ The cyto2 and nuclei models were further trained on the images in this study to achieve high-quality segmentation. Nuclear and/or cellular masks were exported to ImageJ to measure total intensity, mean intensity, and pixel areas of defined regions.

### qPCR

Cells were lysed in TRIzol, and total RNA was isolated using the Direct-zol RNA Miniprep Plus Kit (Zymo #R2072). cDNA was generated using ProtoScript II Reverse Transcriptase (New England Biolabs #M0368L) and poly(dT) primers. 20 μL RT-qPCRs were performed using the PerfeCTa qPCR SuperMix (QuantaBio #95050). Two independent primer sets were used to measure ARH3 mRNA expression: (1) ARH3-Flag primers only amplifying Flag-tagged ARH3 due to a reverse primer spanning the Flag and linker sequence and (2) endo-ARH3 primers amplifying both endogenous and Flag-tagged ARH3 mRNA transcripts (see **Table S2** for primer sequences). To compare mRNA and protein expression levels of ARH3, cells from a single culture were trypsinized and split in half and for either immunoblotting or RNA extraction.

### Cycloheximide chase

U2OS cells were cultured in 6-well plates as described earlier. The half-life of Flag-tagged ARH3 was measured by adding cycloheximide (30 μg/mL, Sigma #C4859) for varying amounts of time, lysing cultures, and immunoblotting for ARH3. ARH3-Flag band intensities were quantified, normalized to Ku70, and expressed relative to DMSO-treated controls. Half-lives were calculated using an exponential decay model in Prism 9 software and statistically compared using 95% confidence intervals (CIs).

### Bacterial protein expression

pDest17 plasmids containing an N-terminal 6x-His tag and Flag-tagged ARH3 wildtype or H182R were transformed into Rosetta (DE3)pLysS Competent Cells (Sigma, #70956). An initial 5 mL culture was grown overnight in Lennox LB broth supplemented with 100 μg/mL of ampicillin and 25 μg/mL of chloramphenicol at 37 °C while shaking at 200 rpm. The following day, cultures were diluted to 0.04 OD600 (1 cm path length) and grown at 37 °C until reaching 0.4 OD600. Proteins were expressed by addition of 1 mM IPTG for 15 hr at 18°C. Cells were harvested by centrifugation, resuspended in ice-cold lysis buffer (25 mM Tris-HCl, pH 8.0, 100 mM NaCl), and sonicated. Following sonication, soluble and insoluble fractions were collected by centrifugation at 20,000xg for 45 min at 4°C.

## RESULTS

### ARH3^H182R^ segregates with a CONDSIAS phenotype in two children

Patient 1 (II-4 in **Figure 1A**) first presented at 28 months old with episodic dystonia provoked by exercise. By age three, he was noted to have developmental delays with especially impacted expressive language. Initial genetic testing for paroxysmal kinesigenic dyskinesias included next generation sequencing of *ADCY5, KCNA1, KCNMA1, PDE10A, PNKD, PRRT2, SLC2A1* and *TRAPPC11* and was negative. Subsequently, an epilepsy panel from an external commercial lab resulted only in variants of uncertain significance in *ARX, CPA6, FASN*, and *ST3GAL3* and was considered negative. Magnetic resonance imaging (MRI) of his brain at 33 months old was also normal/negative (**Figure 1B**). No additional genetic testing was performed until at five years of age when he traveled to Africa with family. Both he and his sister were prescribed atovaquone-proguanil antimalarial prophylaxis for the trip. Unfortunately, Patient 1 died on this trip from neurologic symptoms and respiratory failure in the setting of a diarrheal illness (see **Supplemental Clinical History** for more details).

**Figure 1.**
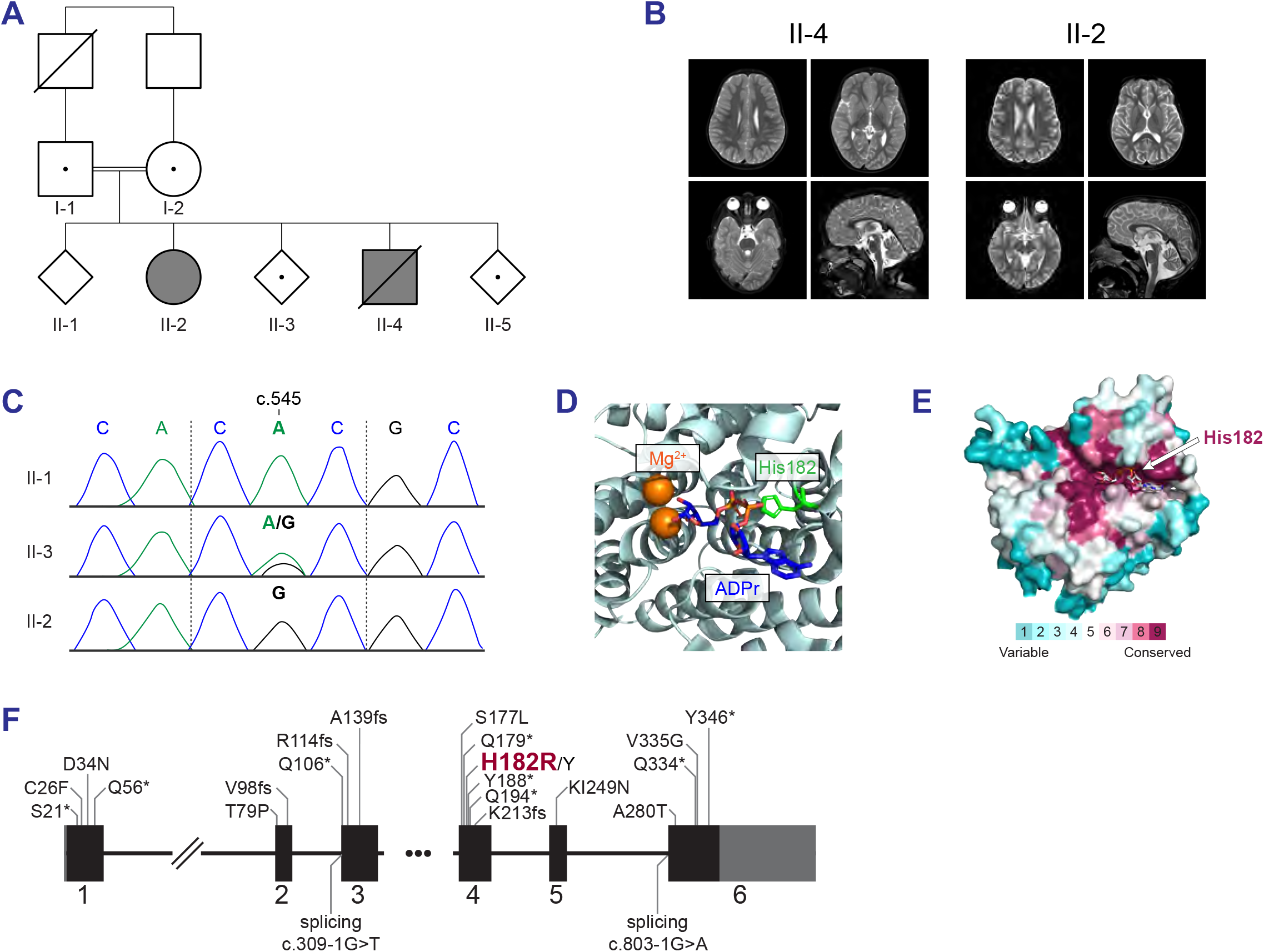
ARH3^H182R^ segregates with CONDSIAS. **(A)** Pedigree analysis of the Somali family in this study. II-2 and II-4, represented with closed symbols, are affected with CONDSIAS and homozygous for ARH3^H182R^. Unaffected individuals heterozygous for ARH3^H182R^ are represented with open symbols and a central dot. Double lines between the parents I-1 and I-2 indicate consanguinity (second cousins). Sex is concealed with diamonds for unaffected siblings to preserve anonymity. **(B)** Normal axial T2 and sagittal T2-FSE brain MRIs of Patient 1 (II-4) at 33 months old and Patient 2 (II-2) at 8 years old. **(C)** Sanger sequencing chromatograms confirming NGS sequencing data. Vertical dashed lines delineate the His182 codon, with CAC encoding histidine and CGC arginine. **(D)** A cartoon model of the ARH3 crystal structure (PDB 6D36) emphasizing the interaction of His182 with ADPr in the active site of ARH3. **(E)** A surface model of the ARH3 crystal structure (PDB 6D36)^39^ colored according to ConSurf conservation scores. **(F)** All ARH3 variants published as occurring in patients with CONDSIAS (see **Table S5** for citations).

Patient 2 (II-2 in **Figure 1A**) is a female of Somali ancestry who presented at age six with epilepsy, intermittent left-sided stiffening, and developmental delays manifesting as difficulty reading, writing, and the need for an individualized education program at school. However, her age of onset is likely younger, as her parents report shaking when sick beginning around 1.5 years old. She also attended the family trip to Africa and took atovaquone-proguanil antimalarial prophylaxis. Like her brother, she developed a diarrheal illness on this trip but survived to present in the United States with subacute, progressive lower extremity weakness, leg pain, and new-onset urinary and fecal incontinence. Brain MRI at this time was negative/normal (**Figure 1B**). Video electroencephalography showed subclinical seizures. She developed respiratory failure and required intubation for diaphragmatic weakness. Electromyogram suggested an acquired, acute motor-predominant peripheral polyneuropathy initially concerning for Guillain-Barré syndrome but with preservation of deep tendon reflexes. Unfortunately, she did not recover independent respiratory function and required tracheostomy for long-term ventilatory support. She has had continued progression with denervation of bilateral lower extremities and distal arms, dyskinesias, and seizures.

A family history revealed unaffected parents with known consanguinity (first cousins) and three additional unaffected children. During the admission of Patient 2, rapid quad-exome analysis of genome sequencing was sent for Patient 2, mother, father and a residual DNA sample from her brother’s (Patient 1’s) prior testing. This testing identified the variants *ADPRS*:NM_017825.2; c.545A>G (p.His182Arg) and *ST3GAL3*:NM_006279.3; c.685G>A (p.Ala229Thr). Both variants were heterozygous in each parent, homozygous in Patients 1 and 2, and determined to be of uncertain significance (**Figures 1A and C**); however, the *ADPRS* homozygous variant was thought to more likely be disease-causing on the basis of clinical phenotype and data suggesting His182 is important for ARH3 function.^28,29^ The *ADPRS* variant calling was confirmed with Sanger sequencing (**Figure 1C**). Targeted testing revealed other siblings as unaffected heterozygotes or non-carriers, indicating the segregation of this *ADPRS* variant with a CONDSIAS phenotype according to a recessive mode of inheritance.

ARH3^H182R^ is predicted to be pathogenic by multiple tools based on evolutionary conservation and protein structure-function relationships (**Table S4**).^30–32^ Further, His182 is a highly evolutionarily conserved residue in the active site of ARH3 that other studies found to be essential for binding ADPr (**Figure 1D-E**).^28,29,33^ ARH3^H182R^ was not found in gnomAD^34^, a collection of sequencing data from over 195,000 individuals taken from many projects, nor in Al Mena^35^, a collection of 2,115 individuals from the Middle East and North Africa. In fact, the only variation at position His182 reported in gnomAD is heterozygous H182Q at a frequency of 2.06e-6, consistent with the autosomal recessive manner of inheritance of CONDSIAS. H182R is also absent from the clinical reporting databases ClinVar and OMIM. However, p.H182Y was identified as a compound heterozygous variant alongside p.Y188* and is reported as likely pathogenic in CONDSIAS, further supporting the pathogenicity of His182 missense variants (**Figure 1F**).^21^

### A model of ARH3 function in U2OS cells

To establish a model of ARH3 function, we first knocked out the *ADPRS* gene in U2OS cells using CRISPR-Cas9 gene editing and a single guide RNA (sgRNA) targeting exon 1 (**Figure 2A**). As expected, ARH3 was not detected in ARH3^-/-^ (ARH3^KO^) cells (**Figure 2B)**. Additionally, ARH3^KO^ cells exhibited no cell cycle defects (**Figure S1A-C**). Since ARH3 is the only mono-ADP-ribosylhydrolase known to remove serine-linked mono(ADP-ribose) (Ser-MAR), we monitored MARylated protein levels in ARH3^WT^ and ARH3^KO^ total cell extracts using two antibodies specific for pan-MAR (MAR, AbD33204) and serine-specific MAR (Ser-MAR, AbD33205). ARH3^KO^ cells exhibited increased protein MARylation compared to wildtype cells. Importantly, overexpression of exogenous Flag-tagged ARH3 (ARH3^WT^-Flag) and/or addition of a PARP1/2-inhibitor (PARPi, Olaparib) completely reserved the elevated protein MARylation seen in the ARH3^KO^ cells to levels observed in wildtype cells (**Figures 2C-D**). Interestingly, comparison of ARH3^WT^ and ARH3^KO^ cells revealed three additional lower molecular weight bands between 23 to 33 kDa specific to ARH3.

**Figure 2.**
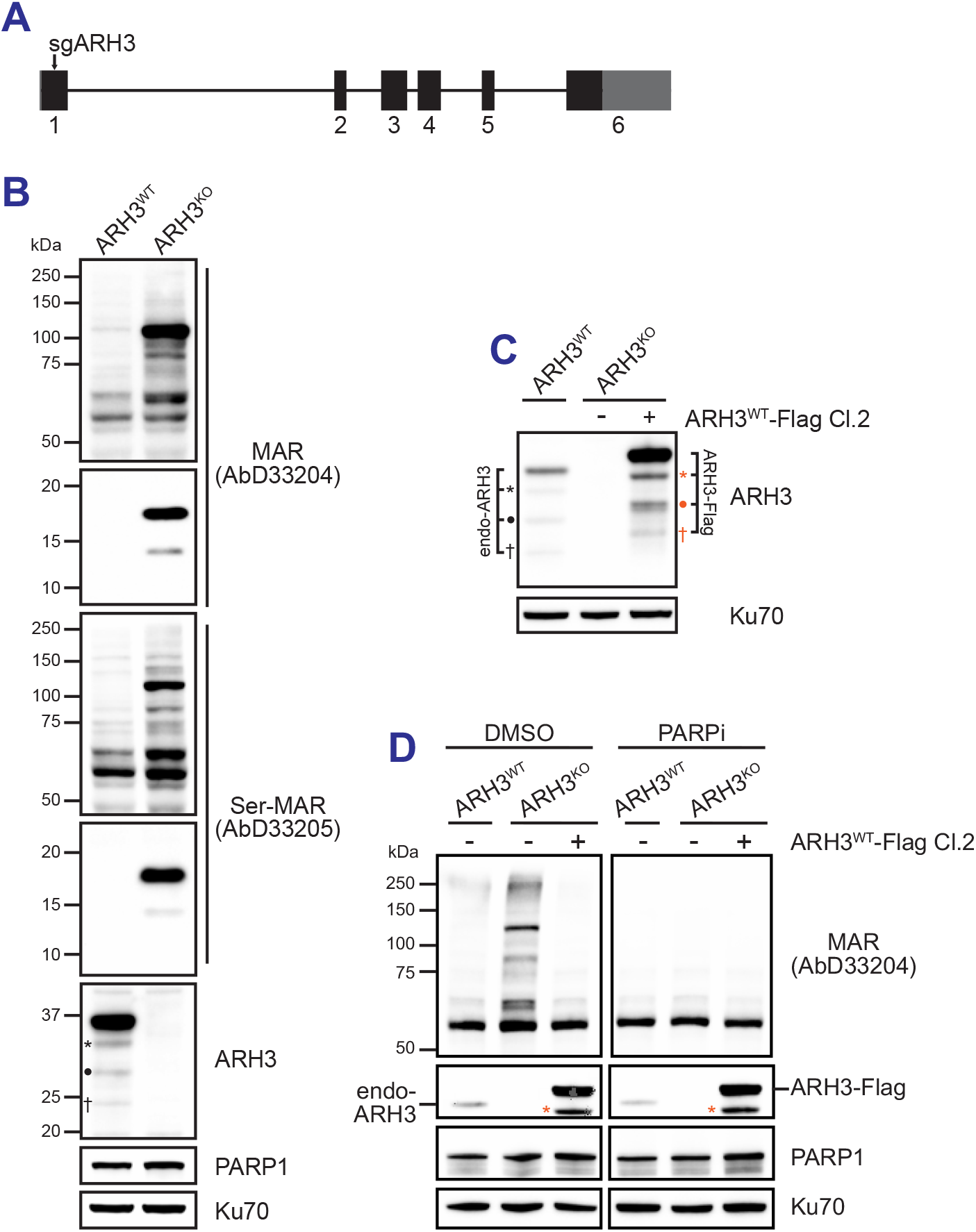
Establishing a cellular model of ARH3 function. **(A)** Cartoon depiction of the CRISPR/Cas9 sgRNA targeting site along the *ADPRS* gene locus. Boxes represent exons that are numerically indexed below; lines represent introns. 5’- and 3’-UTRs are gray, and coding regions are black. **(B)** Immunoblot analysis of ARH3 protein expression and proteins modified by MAR or Ser-MAR in whole cell lysates of U2OS ARH3^WT^ and ARH3^KO^ cells. The *, ·, and † denote putative, endogenous ARH3 isoforms that disappeared in ARH3^KO^ cells. **(C)** Immunoblot analysis of ARH3 protein expression in U2OS ARH3^WT^ cells, ARH3^KO^ cells, and a single clone of ARH3^KO^ cells stably expressing Flag-tagged ARH3^WT^. Note, the molecular weight of different ARH3^WT^-Flag isoforms is due to the addition of the Flag tag. **(D)** Immunoblot analysis of ARH3, PARP1, and MARylated proteins in U2OS ARH3^WT^ cells, ARH3^KO^ cells, and a single clone of ARH3^KO^ cells stably expressing Flag-tagged ARH3^WT^ treated with either DMSO or 10 μM Olaparib (PARPi) for 6 hr. The asterisk is correlating the band here with those in (B) and (C).

Overexpression of exogenous Flag-tagged ARH3 (ARH3^WT^-Flag) also confirmed different protein isoforms of ARH3 (**Figures 2B-C**). Given that only a single mRNA isoform has been detected for ARH3, these data suggest that either multiple ARH3 protein isoforms are translated from the same mRNA or that ARH3 is post-translationally cleaved.

### The H182R variant disrupts expression and nuclear localization of ARH3

To assess the impact of the H182R variant on ARH3 function, we next complemented ARH3^KO^ cells with ARH3^H182R^-Flag. We noticed that the ARH3^H182R^-Flag expression is significantly lower than ARH3^WT^-Flag in multiple independent clones (**Figure 3A**). Further, the decreased ARH3^H182R^-Flag protein level was not due to lower ARH3^H182R^-Flag mRNA expression measured by qPCR using independent primer sets targeting either ARH3 or ARH3-Flag (**Figures 3B-D**). Consistent with the immunoblot analysis, total ARH3^H182R^-Flag protein measured by Flag immunofluorescence microscopy is lower than ARH3^WT^-Flag (**Figure 3E-F**). Furthermore, whereas ARH3^WT^-Flag localizes to both nuclear and cytoplasmic regions, the residual ARH3^H182R^-Flag is almost entirely cytoplasmic (**Figures 3E, G-H**). The nuclear-to-cytoplasmic ratio of ARH3^H182R^-Flag expression is significantly lower compared to ARH3^WT^-Flag expression (**Figure 3I**). Taken together, these data show that ARH3^H182R^ is poorly expressed in cells and that even the reduced amount of mutant protein fails to localize to the nucleus.

**Figure 3.**
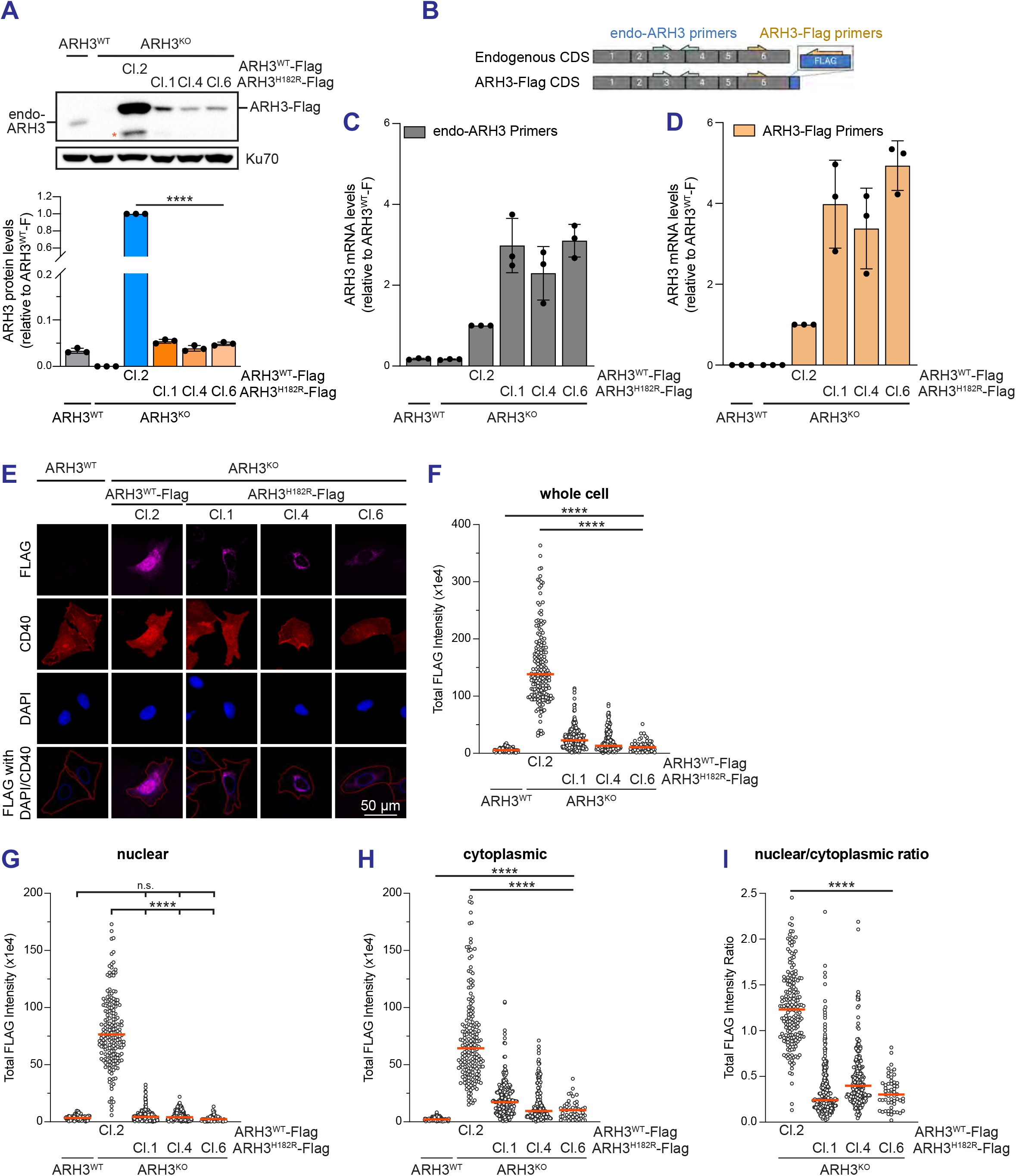
ARH3^H182R^ is poorly expressed in cells and loses nuclear localization. **(A)** Immunoblot analysis of ARH3 protein expression in U2OS ARH3^WT^, ARH3^KO^ cells, and ARH3^KO^ cells stably expressing either ARH3^WT^-Flag or ARH3^H182R^-Flag is shown with an asterisk denoting the same band seen in Figure 2. Total band intensity was quantified, normalized to Ku70, and expressed relative to ARH3^WT^-Flag. Statistical analysis used a one-way ANOVA followed by Tukey’s multiple comparisons test (****, p<0.0001). **(B-D)** RT-qPCR analysis of ARH3 mRNA expression in indicated cell lines using primers targeting both endogenous and Flag-tagged ARH3 (endo-ARH3 primers) (C) or primers specific to Flag-tagged ARH3 (ARH3-Flag primers) (D). **(E-I)** The subcellular localization of ARH3^WT^-Flag and ARH3^H182R^-Flag was evaluated by immunofluorescence using a monoclonal Flag antibody. Representative images of cells probed for Flag (ARH3), CD40 (plasma membrane), and DAPI (nuclei) are shown in (E). Segmentation masks for nuclei and cells are shown in blue and red, respectively. Total ARH3-Flag fluorescence intensity of whole cells (F), individual nuclei (G), cytoplasmic regions (H), and the ratio of nuclear to cytoplasmic intensity in each cell (I) were quantified (n > 220 per sample). Orange bars indicate medians. Statistical analyses used a nonparametric one-way ANOVA followed by Dunn’s multiple comparisons test (****, p<0.0001; n.s., not significant).

### The H182R variant destabilizes the ARH3 protein in cells

The strikingly diminished steady-state abundance of the ARH3^H182R^ protein prompted us to investigate the kinetics of ARH3^WT^ and ARH3^H182R^ degradation in cells. We treated AHR3^WT^ and ARH3^H182R^ cells with cycloheximide (CHX), an inhibitor of protein synthesis, and monitored protein abundance over 8 hours. While ARH3^WT^-Flag abundance was unchanged over 8 hours in CHX, ARH3^H182R^-Flag levels dramatically decreased with a half-life of 2.4 hours (**Figures 4A-B**). The reduced ARH3^H182R^ stability was observed in three independent clones treated with CHX for 8 hours (**Figures 4C-D**). To corroborate the cycloheximide chase experiment, we expressed His-tagged ARH3^WT^ and ARH3^H182R^ in *E. coli* to measure their relative solubility following cell lysis. After ARH3 protein expression was induced with IPTG, cell lysates were centrifuged to recover into soluble and insoluble fractions. The expression of both ARH3 wildtype and H182R were similar in whole cell lysates (**Figure 4E**, left panel). However, ARH3^WT^ was predominantly in the soluble fraction and ARH3^H182R^ mainly in the insoluble fraction (**Figure 4E**, right panel). Similar insolubility was observed using GST-tagged ARH3^H182R^ in *E. coli* (*data not shown*). Together, these results indicate that the H182R variant negatively impacts ARH3 protein stability, and therefore its proper enzymatic functioning, to overall increase MAR levels in cells to the same extent as ARH3^KO^ cells.

**Figure 4.**
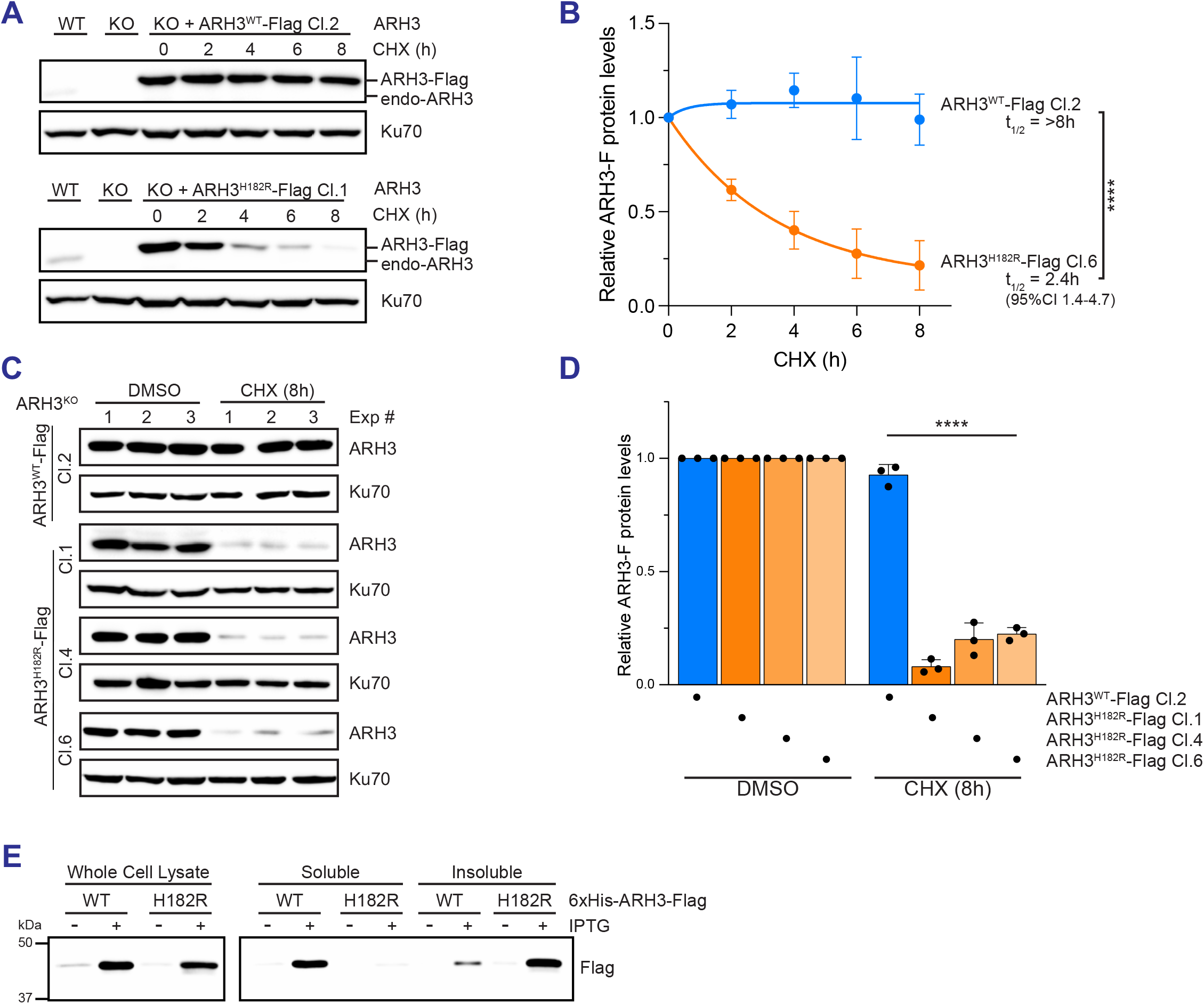
The H182R variant destabilizes ARH3 protein. **(A-B)** Single clones of ARH3^KO^ cells stably expressing either ARH3^WT^-Flag or ARH3^H182R^-Flag were treated with cycloheximide (CHX, 30 μg/mL) and collected at the indicated time points (n=3). Representative immunoblot analyses are shown in (A). ARH3-Flag band intensity was quantified, normalized to Ku70, and expressed relative to DMSO-treated controls. The half-lives of ARH3^WT^-Flag and ARH3^H182R^-Flag were calculated from an exponential decay model of the data in (B). Statistical analysis was performed using the 95% CI for the half-life ARH3^H182R^-Flag against a reference value of 8 hours (****, p<0.0001). **(C-D)** Single clones of ARH3^KO^ cells stably expressing either ARH3^WT^-Flag or ARH3^H182R^-Flag were treated with cycloheximide (CHX, 30 μg/mL) for 8 hrs. ARH3-Flag band intensity was quantified from the immunoblots in (C), normalized to Ku70, and expressed relative to DMSO-treated controls in (D). **(E)** 6xHis- and Flag-tagged ARH3^WT^ and ARH3^H182R^ proteins were expressed in *E. coli* by the addition of IPTG. Expression and solubility of ARH3 wildtype and mutants were monitored in whole cell lysates, soluble, and insoluble fractions using anti-Flag.

### The ARH3^H182R^ mutant fails to suppress mono(ADP-ribosylation) in ARH3^KO^ cells

The combination of failed ARH3^H182R^ nuclear localization with poor expression secondary to destabilization and rapid degradation suggests that ARH3 function and subsequent ADP-ribosylation dynamics are impaired in ARH3^H182R^-expressing cells. Indeed, expressing ARH3^H182R^-Flag failed to rescue the elevated protein MARylation seen in ARH3^KO^ cells, while ARH3^WT^-Flag expression completely reversed the MARylation to levels observed in wildtype cells (**Figure 5A**). Further, the low ARH3^H182R^ expression in the nucleus correlated with increased nuclear MAR in cells expressing ARH3^H182R^-Flag compared to ARH3^WT^-Flag (**Figures 5B-C**). Taken together, ARH3^H182R^ appears to functionally mimic the ARH3^KO^ cells as a consequence of the observed defects in expression, stability, and localization.

**Figure 5.**
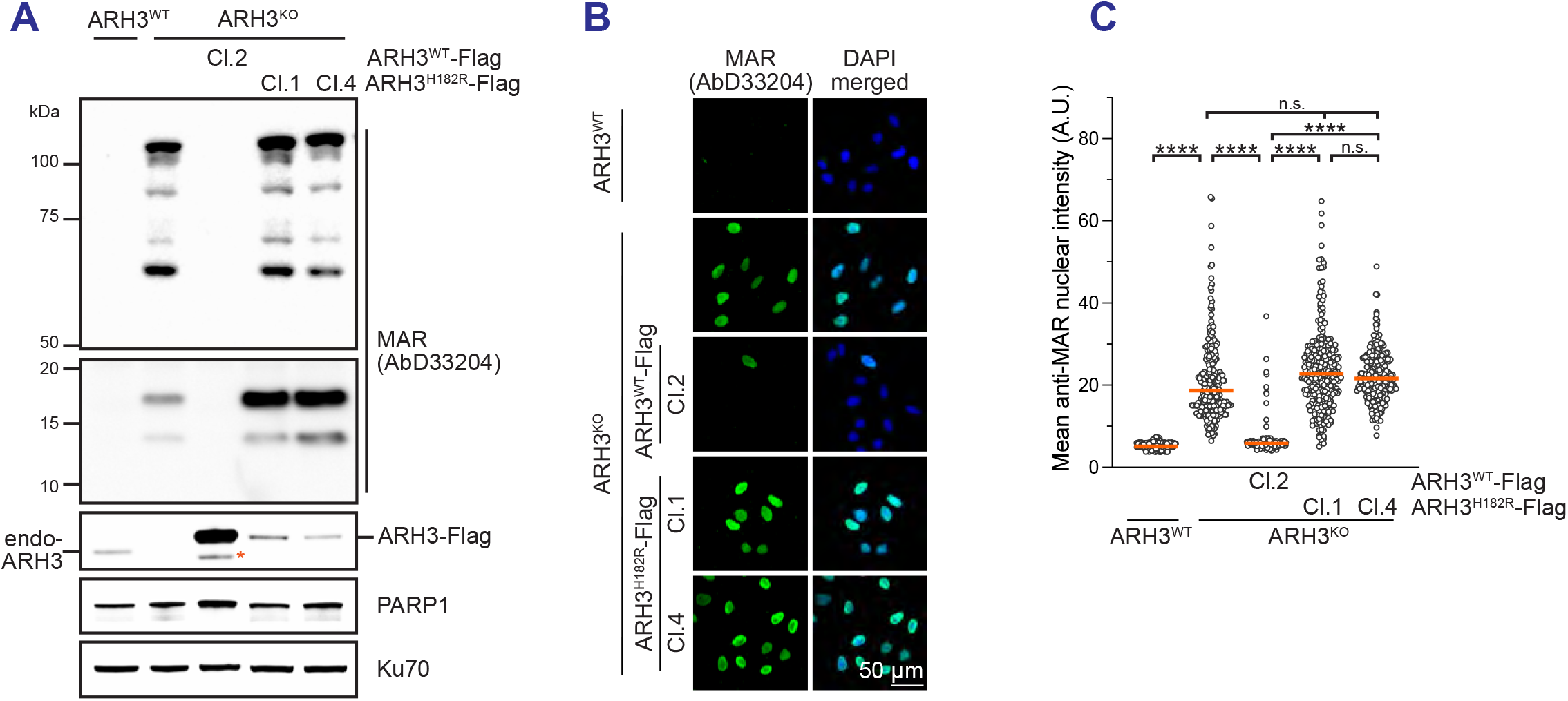
ARH3^H182R^ fails to resolve MARylation in ARH3^KO^ cells. **(A)** Immunoblot analysis of ARH3, PARP1, and proteins modified by MAR in whole cell lysates of U2OS ARH3^WT^, ARH3^KO^ cells, and ARH3^KO^ cells stably expressing either ARH3^WT^-Flag or ARH3^H182R^-Flag. The orange asterisk denotes the same band seen in Figure 2B. **(B-C)** Immunofluorescence analysis of nuclear MAR in pre-extracted U2OS ARH3^WT^ cells, pooled ARH3^KO^ cells, and single clones of ARH3^KO^ cells stably expressing Flag-tagged ARH3^WT^ or ARH3^H182R^. Representative images are shown in (B) and MAR intensity of 275 individual nuclei with orange bars at the median in (C). All statistical analyses used a nonparametric one-way ANOVA followed by Dunn’s multiple comparisons test (****, p<0.0001; n.s., not significant).

## DISCUSSION

In this study, we provided a detailed biochemical characterization of a novel loss-of-function variant in *ADPRS* in two patients with CONDSIAS. The ARH3^H182R^ variant fails to localize to the nucleus, is structurally destabilized, and is rapidly degraded in cells, resulting in the inability to resolve accumulated MARylated proteins in ARH3^KO^ cells. Our clinical report marks cases 50 and 51 of CONDSIAS reported in the literature to date (**Table S5**), representing 29 families and 23 variants (**Figure 1F**). Although these case reports are essential to developing our understanding of CONDSIAS, functional characterization of the many different *ADPRS* variants is needed to deduce pathogenic mechanisms of CONDSIAS.

A comprehensive mechanism explaining the severe neurodegeneration following loss of ARH3 activity remains elusive. While many patients unfortunately pass away between the ages of 5-10, patients identified with the ARH3^V335G^ variant in six different families tend to live substantially longer, even into their 30’s and 50’s (**Figure S2A**). This is likely because ARH3^V335G^ retains some enzymatic activity but is expressed at a lower level and has altered subcellular localization.^19^ In particular, ARH3^V335G^ is almost exclusively found in the nucleus and mitochondria but not the cytoplasm. These favorable clinical phenotypes associated with the V335G variant may suggest that the primary dysfunction caused by ARH3 deficiency is associated with ARH3 activity in the nucleus. Similar to the ARH3^V335G^ variant, the H182R variant found in our patients exhibited reduced protein expression but mainly localized to the cytoplasm, prohibiting the ARH3 activity in the nucleus. Although we could not directly determine the activity of residual ARH3^H182R^ in cells, it is likely that the disruption of ADPr binding caused by mutation of His182 is detrimental to enzymatic activity. Another variant, ARH3^A280T^, identified in four patients also seems to correlate with longer life, but nothing is known about its subcellular localization and expression.^20^ Future experiments are necessary to characterize the effect of other pathogenic variants on ARH3 protein stability, localization, and enzymatic activities.

Many aspects of the clinical presentation of CONDSIAS require better models to investigate. For example, why do children develop normally early in life and then deteriorate around two years of age (**Figure S2A**)? This presentation is consistent with a progressive disease caused by accumulation of cellular insults over time, but it is unclear if loss of ARH3 is differentially harmful *in utero* compared to after birth. A very similar parallel exists with the specificity of toxicity of ARH3 deficiency for neuronal tissue: what are the factors that make neurons sensitive while other systems (e.g., hematopoietic) seem intact? Unfortunately, no suitable animal models have been developed to address these questions: ARH3^KO^ mice fail to recapitulate the phenotype observed in humans^14^, and in *Drosophila* there is no ARH3 homolog (*Drosophila* PARG can remove serine-MAR).^36^

Further, the episodic nature of CONDSIAS being seemingly linked to physiologic stressors has to date been poorly modeled *in vitro*, with most studies using hydrogen peroxide as a rather blunt insult. The most common physiologic challenge reported to precipitate severe episodes in children with CONDSIAS is febrile illness which could be modeled as inflammatory processes in cells. The children in this study, in addition to developing a diarrheal illness, were also exposed to atovaquone-proguanil antimalarial prophylaxis which could increase reactive oxygen species (ROS).^37^ It is possible that the combination of atovaquone-proguanil and febrile illness made these children even more sensitive to ARH3 deficiency, explaining the severity of this particular episode that unfortunately resulted in the death of Patient 1 and substantial decompensation of Patient 2. In addition, recent work reported that RNA can be modified by ADPr in response to inflammatory signaling, and that ARH3 is able to remove this modification.^9^ Investigating these and other mechanisms of neurodegeneration driven by ARH3 deficiency in CONDSIAS may offer unique insights into the pathologies of other neurodegenerative diseases with dysregulated ADPr homeostasis, like amyotrophic lateral sclerosis (ALS), Parkinson’s disease, Alzheimer’s disease, Huntington’s disease, and multiple sclerosis.^38^

In summary, our clinical case report adds to the critical information on the disease course and management of CONDSIAS, and our experimental results provide a detailed biochemical characterization of a loss-of-function variant in ARH3 never before identified in patients. Our functional data, combined with the clinical presentation of the children, *in silico* predictions of H182R effect on ARH3, and population genetics data, update the ARH3^H182R^ variant from one of uncertain significance to pathogenic according to the criteria laid out by the ACMG and AMP (**Table S6**). Finally, these results add to a growing body of evidence that ADP-ribosylation has central roles in neuronal development and function.

## Supporting information

Supplemental Clinical Histories

## Figure Legends

**Supplementary Figure 1.**
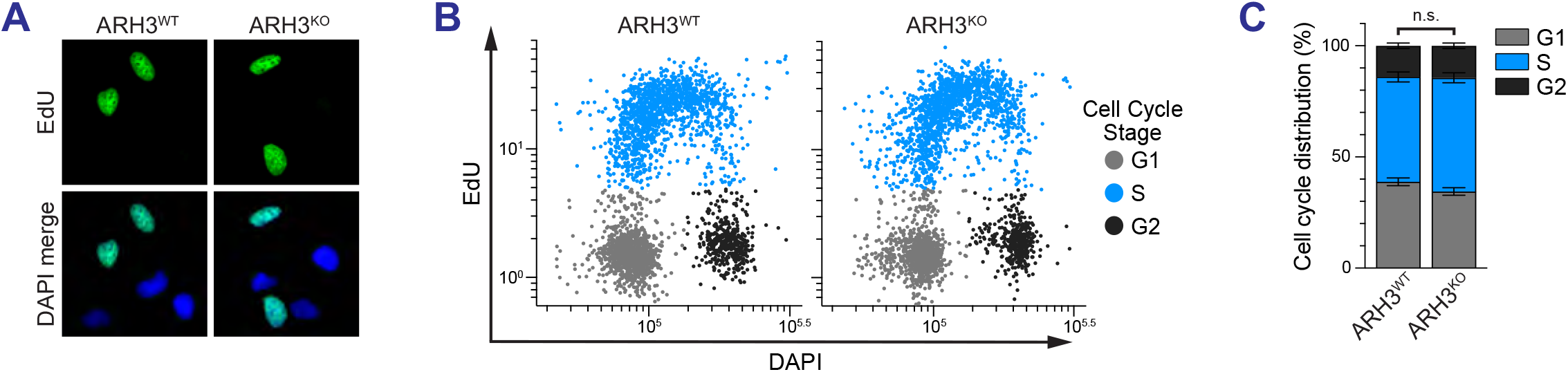
**(A-C)** Representative images of ARH3^WT^ and ARH3^KO^ cells stained for EdU and DAPI are shown in (A). Quantitative image-based cytometry showing the cell cycle distribution of individual ARH3^WT^ and ARH3^KO^ cells based on total DNA content (DAPI) and DNA synthesis measured by EdU incorporation is shown in (B). Frequencies of G1, S, and G2 cell-cycle stages are plotted in C. Statistical analysis used a one-way ANOVA followed by Tukey’s multiple comparisons test (n.s., not significant).

**Supplementary Figure 2.**
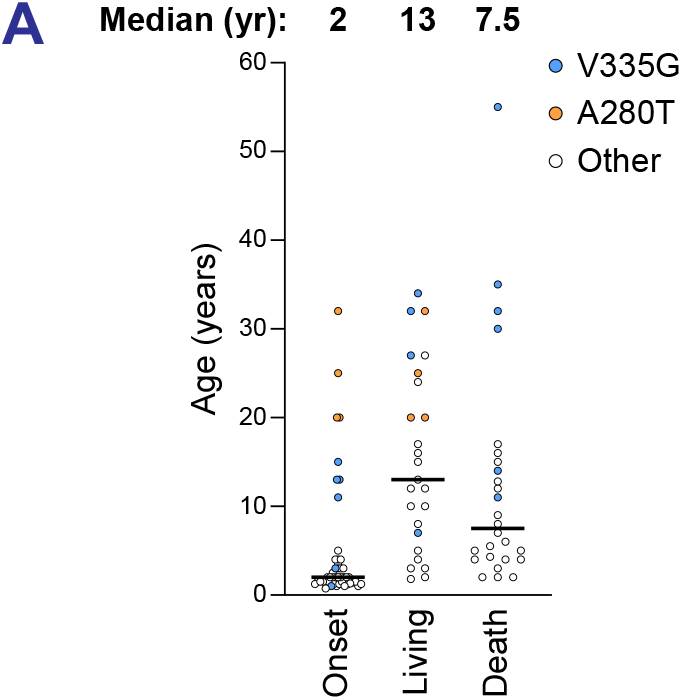
**(A)** Age of disease onset, death, and most recent age of those living at time of publication for all patients reviewed in **Table S5**.

## Data Availability

Data collected and analyzed for this manuscript are available upon request.

## Acknowledgements

We are grateful to the patients and their families for participation in this study, and to all medical and laboratory specialists for providing clinical care. We thank Dr. Hibbs for providing genetic counseling with the family.

H.D. Nguyen was supported by grants from the Edward P. Evans Foundation, American Society of Hematology, the NIH’s National Center for Advancing Translational Sciences, grants KL2TR002492 and UL1TR002494, the National Heart, Lung, and Blood Institute (R01 HL163011), and the 2022 AACR Career Development Award to Further Diversity, Equity, and Inclusion in Cancer Research, which is supported by Merck, grant number 22–20–68-NGUY.

## Author Contributions

Conceptualization: C.B.Jr, H.D.N, Data curation: M.B., S.B., A.A., K.H., Formal analysis: M.B., S.B., C.B.Jr., H.D.N., Writing-original draft: M.B., C.B,Jr., H.D.N., Writing-review & editing: M.B., S.B., C.B. Jr., H.D.N.

## Ethics Declaration

The University of Minnesota Institutional Review Board (IRB) determined that the human subjects component of this work was not human subjects research (reviewed as STUDY00021345).

