## Supplemental Tables for "A novel variant in *ADPRS* disrupts ARH3 stability and subcellular localization in children with neurodegeneration and respiratory failure"

**Supplemental Table 1: List of antibodies used in this study**

| Antibodies | Clone | Host | Company | Cat # | Dilution |
|---|---|---|---|---|---|
| ADPRHL2 (ARH3) | polyclonal | rabbit | Millipore Sigma | HPA027104 | 1:1000 |
| MAR | AbD33204 | human/rabbit chimera | Bio-Rad | HCA354 | 1:500 |
| Ser-MAR | AbD33205 | human/rabbit chimera | Bio-Rad | HCA355 | 1:500 |
| PARP1 | 46D11 | rabbit | CST | 9532S | 1:1000 |
| Ku70 | 1.5 | mouse | GeneTex | GTX70271 | 1:5000 |
| FLAG | M2 | mouse | Sigma-Aldrich | F1804 | 1:1000 |
| FLAG | D6W5B | rabbit | CST | 14793S | 1:1000 |
| CD40 | EPR20735 | rabbit | abcam | ab224639 | 1:500 |

**Supplemental Table 2: List of primers and sequences used in this study**

| Primers | Sequence | Gene/Assay |
|---|---|---|
| MBp085_ARH3_F | AACGGGAAAGGCTCCTATGG | *ADPRS* (ARH3) Expression. "endo-ARH3 primers" |
| MBp086_ARH3_R | CGGGCAAACTTCTGCACATC | |
| MBp087_exoARH3_F | ATTGCTGGTGCCTACTATGGG | *ADPRS-Flag* (ARH3-FLAG) Expression. "ARH3-Flag primers" |
| MBp088_exoARH3_R | TCGTCTTTGTAGTCGGTGGCG | |
| VCp009_GAPDH_F | GACATCAAGAAGGTGGTGAAGCAG | *GAPDH* Expression |
| VCp010_GAPDH_R | AGCTTGACAAAGTGGTCGTTGAG | |
| ADPRS_Seq_c545A>G_F | GAACCCCAAATGTCGCGATG | *ADPRS* NM_017825.3 c.545 sequencing |
| ADPRS_Seq_c545A>G_R | CCACATACCCACACCACACA | |

| sgRNA | Sequence | Gene |
|---|---|---|
| sgARH3-2 | GCGCTGCTCGGGGACTGCGT | *ADPRS* (ARH3) |

**Supplemental Table 3: List of plasmids used in this study**

| Plasmid Name | Source | Cat # | Comments |
|---|---|---|---|
| pSpCas9(BB)-2A-Puro (PX459) V2.0 | Addgene | 62988 | received from Dr. Clark Chen at the University of Minnesota |
| pLenti6.2-ccdB-3xFLAG-V5 | Addgene | 87071 | |
| attb1-ARH3-Flag-attb2 | GeneUniversal | | synthesized ARH3 WT and H182R constructs |
| pCMV-dR8.2 dvpr | Addgene | 8455 | |
| pCMV-VSV-G | Addgene | 8454 | |
| Gateway pDEST17 | Invitrogen | 11803012 | |

**Supplemental Table 4:** ***in silico*** **pathogenicity predictions**

| ARH3H182R | | MutationTaster† | CADD‡ | REVEL* |
|---|---|---|---|---|
| Coding change (NM_017825.3) | c.545A<G | Disease Causing | 26.3 | 0.759 |
| Protein change (NP_060295.1) | p.H182R | | | |
| Genomic change (GRCh38/hg38) | chr1:36091938A>G | | | |

† Predicted "disease causing" with a Bayesian Classifier Probability of 0.999999922463634

‡ CADD v1.6, scores are ranked relative to all possible substitutions of the human genome such that scores ≥20 indicate the top 1% of most deleterious SNVs and ≥30 indicate the 0.1% most deleterious.

* REVEL is an ensemble prediction algorithm of 13 different tools. This REVEL score correlates to a specificity of 0.969, meaning that only ~3% of neutral variants are expected to have a REVEL score this high or higher

**Table S5 Sheet 1: CONDSIAS Publications**

| PMID | First Author | Publication Year | Journal | Title | DOI |
|---|---|---|---|---|---|
| 37712079 | Mahungu AC | 2023 | Front Neurol | The mutational profile in a South African cohort with inherited neuropathies and spastic paraplegia | 10.3389/fneur.2023.1239725 |
| 37392332 | Lindskov FO | 2023 | Cerebellum | Expanding the Spectrum of Stress-Induced Childhood-Onset Neurodegeneration with Variable Ataxia and Seizures (CONDSIAS) | 10.1007/s12311-023-01582-w |
| 36911439 | Bajaj S | 2022 | Ann Indian Acad Neurol | An Indian Child with CONDSIAS Due to a Novel Variant in ADPRHL2 Gene | 10.4103/aian.aian_558_22 |
| 35693655 | Ozturk G | 2022 | Ann Indian Acad Neurol | Stress-induced Childhood Onset Neurodegeneration with Ataxia and Seizures (CONDSIAS) Presenting with Torticollis Attacks: Phenotypic Variability of the Same Mutation in Two Turkish Patients | 10.4103/aian.aian_314_21 |
| 35664652 | Ma J | 2022 | Front Genet | Child-Onset Cerebellar Ataxia Caused by Two Compound Heterozygous Variants in ADPRS Gene: A Case Report | 10.3389/fgene.2021.788702 |
| 35222245 | Lu A | 2022 | Front Neurol | Case Report: Stress-Induced Childhood-Onset Neurodegeneration With Ataxia-Seizures Syndrome Caused by a Novel Compound Heterozygous Mutation in ADPRHL2 | 10.3389/fneur.2022.807291 |
| 34479984 | Beijer D | 2021 | Life Sci Alliance | Biallelic ADPRHL2 mutations in complex neuropathy affect ADP ribosylation and DNA damage response | 10.26508/lsa.202101057 |
| 33528672 | Durmus H | 2021 | Neurol Sci | Episodic psychosis, ataxia, motor neuropathy with pyramidal signs (PAMP syndrome) caused by a novel mutation in ADPRHL2 (AHR3) | 10.1007/s10072-021-05100-w |
| 33426173 | Mishra B | 2020 | Mov Disord Clin Pract | Dystonia and Myelopathy in a Case of Stress-Induced Childhood-Onset Neurodegeneration with Ataxia and Seizures (CONDSIAS) | 10.1002/mdc3.13125 |
| 32746785 | Aryan H | 2020 | BMC Neurol | Novel imaging and clinical phenotypes of CONDSIAS disorder caused by a homozygous frameshift variant of ADPRHL2: a case report | 10.1186/s12883-020-01873-3 |
| 30830864 | Mashimo M | 2019 | JCI Insight | PARP1 inhibition alleviates injury in ARH3-deficient mice and human cells | 10.1172/jci.insight.124519 |
| 30401461 | Danhauser K | 2018 | Am J Hum Genet | Bi-allelic ADPRHL2 Mutations Cause Neurodegeneration with Developmental Delay, Ataxia, and Axonal Neuropathy | 10.1016/j.ajhg.2018.10.005 |
| 30100084 | Ghosh SG | 2018 | Am J Hum Genet | Biallelic Mutations in ADPRHL2, Encoding ADP-Ribosylhydrolase 3, Lead to a Degenerative Pediatric Stress-Induced Epileptic Ataxia Syndrome | 10.1016/j.ajhg.2018.07.010 |

**Table S5 Sheet 2: CONDSIAS Patients**

| PMID | First Author | Publication Year | Family Count | Patient Count | Variant Coding | Variant Protein | Onset of symptoms | Age Living | Age Death | Notes |
|---|---|---|---|---|---|---|---|---|---|---|
| 30100084 | Ghosh SG | 2018 | Family 1 | Patient 1 | c.1000C>T | Q334* | 2 | | 15 | |
| | | | | Patient 2 | c.1000C>T | Q334* | 1.5 | | 9 | |
| | | | | Patient 3 | c.1000C>T | Q334* | 1.5 | | 7 | |
| | | | | Patient 4 | c.1000C>T | Q334* | 1.5 | | 4 | |
| | | | | Patient 5 | c.1000C>T | Q334* | 1.5 | | 2 | |
| | | | | Patient 6 | c.1000C>T | Q334* | 2 | | 2 | |
| | | | | Patient 7 | c.1000C>T | Q334* | 1.25 | | 2 | |
| | | | | Patient 8 | c.1000C>T | Q334* | 1.25 | 4 | | |
| | | | | Patient 9 | c.1000C>T | Q334* | 1.3 | 3 | | |
| | | | Family 2 | Patient 10 | c.316C>T | Q106* | 2 | 16 | | |
| | | | Family 3 | Patient 11 | c.235A>C | T79P | 4 | 15 | | |
| | | | Family 4 | Patient 12 | c.414_418del | A139fs | 2 | 13 | | |
| | | | | Patient 13 | c.414_418del | A139fs | 0.75 | 2 | | |
| | | | Family 5 | Patient 14 | c.530C>T | S177L | 1 | | 6 | |
| | | | | Patient 15 | c.530C>T | S177L | 1.5 | 3 | | |
| | | | Family 6 | Patient 16 | c.100G>A | D34N | | 10 | | |
| 30401461 | Danhauser K | 2018 | Family 7 | Patient 17 | c.1004T>G | V335G | 1.4 | | 14 | |
| | | | | Patient 18 | c.1004T>G | V335G | 1 | 27 | | |
| | | | Family 8 | Patient 19 | c.744_746del | KI249N | 4 | | 17 | |
| | | | Family 9 | Patient 20 | c.1038C>G | Y346* | 2 | 12 | | |
| | | | Family 10 | Patient 21 | c.1004T>G | V335G | 11 | | 30 | |
| | | | | Patient 22 | c.1004T>G | V335G | 13 | 32 | | |
| | | | Family 11 | Patient 23 | c.1004T>G | V335G | 3 | 7 | | |
| | | | Family 12 | Patient 24 | c.1004T>G | V335G | 2 | | 11 | |
| | | | Family 13 | Patient 25 | c.309-1G>T | splicing | 1 | | 12.8 | |
| | | | | Patient 26 | c.309-1G>T | splicing | 2 | | 5 | |
| | | | Family 14 | Patient 27 | c.292del | V98fs | 1.25 | | 4.3 | |
| | | | | Patient 28 | c.292del | V98fs | 1.2 | 1.8 | | |
| 30830864 | Mashimo M | 2019 | Family 15 | Patient 29 | c.340_341del | R114fs | | | 8 | Assumed affected based on clinical history. |
| | | | | Patient 30 | c.340_341del | R114fs | | | 4 | Assumed affected based on clinical history. |
| | | | | Patient 31 | c.340_341del | R114fs | 1.5 | 8 | | |
| | | | | Patient 32 | c.340_341del | R114fs | | 27 | | |
| 32746785 | Aryan H | 2020 | Family 16 | Patient 33 | c.636_639del | K213fs | 4 | | 4 | |
| 33426173 | Mishra B | 2020 | Family 17 | Patient 34 | c.100G>A | D34N | 3 | 12 | | |
| 33528672 | Durmus H | 2021 | Family 18 | Patient 35 | c.838G>A | A280T | 32 | 32 | | |
| | | | | Patient 36 | c.838G>A | A280T | 25 | 25 | | |
| | | | | Patient 37 | c.838G>A | A280T | 20 | 20 | | |
| | | | Family 19 | Patient 38 | c.838G>A | A280T | 20 | 20 | | |
| 34479984 | Beijer D | 2021 | Family 20 | Patient 39 | c.1004T>G | V335G | 15 | | 32 | |
| | | | | Patient 40 | c.1004T>G | V335G | 13 | 34 | | |
| | | | Family 21 | Patient 41 | c.77G>T | C26F | 1.25 | | 16 | |
| 35222245 | Lu A | 2022 | Family 22 | Patient 42 | c.62C>A; c.535C>T | S21*; Q179* | 3 | 17 | | |
| 35664652 | Ma J | 2022 | Family 23 | Patient 43 | c.508C<T; c.803-1G>A | Q194*; splicing | 2.5 | | 3 | |
| 35693655 | Ozturk G | 2022 | Family 24 | Patient 44 | c.235A>C | T79P | 1 | | 5.5 | |
| | | | Family 25 | Patient 45 | c.235A>C | T79P | 3.5 | 5 | | |
| 36911439 | Bajaj S | 2022 | Family 26 | Patient 46 | c.166C>T | Q56* | 2 | | 12 | |
| 37392332 | Lindskov FO | 2023 | Family 27 | Patient 47 | c.544C>T; c.564C>A | H182Y; Y188* | 5 | 24 | | Currently treating with minocycline as an alternative PARP1 inhibitor |
| 37712079 | Mahungu AC | 2023 | Family 28 | Patient 48 | c.1004T>G | V335G | | | 55 | |
| | | | | Patient 49 | c.1004T>G | V335G | | | 35 | Assumed affected based on clinical history. Died in 30's secondary to a fall. |
| This study | Bannister M | 2024 | Family 29 | Patient 50 | c.545A>G | H182R | 2 | | 5 | |
| | | | | Patient 51 | c.545A>G | H182R | 3 | 11 | | |
