## Supplemental Clinical Histories for "A novel variant in *ADPRS* disrupts ARH3 stability and subcellular localization in children with neurodegeneration and respiratory failure"

### Patient 1

Patient 1 (II-4 in **Figure 1A**) is a male of Somali ancestry born in the United State and is brother and second cousin to patient 2 (II-2 in **Figure 1A**). There were no complications with pregnancy nor his birth. He was walking at approximately one year of age, but speech was delayed. Beginning at 28 months of age, he had dystonic episodes lasting 20-30 minutes with left-sided stiffness in the lower extremities and loss of control of upper extremities. These episodes seemed to be induced by activity. He remained conscious and able to communicate without deviation of the eyes nor change in facial expression during these episodes. Initial evaluation otherwise revealed normal gross and fine motor movement.

He presented to neurology at 32 months of age when genetic testing for paroxysmal kinesigenic dyskinesias was performed as a slice from exome sequencing for *ADCY5*, *KCNA1*, *KCNMA1*, *PDE10A*, *PNKD*, *PRRT2*, *SLC2A1* and *TRAPPC11.* This initial genetic testing resulted as negative. A magnetic resonance imaging (MRI) of his brain done at 33 months was also normal/negative (**Figure 1B**). At 33 months of age, an electroencephalography (EEG) was abnormal due to the presence of low-amplitude spikes in the right posterior temporal region primarily during drowsiness. The EEG background was normal for his age. He was assessed at 34 months as having temporal lobe epilepsy with exercise induced seizures, and he was started on oxcarbazepine at 17.5mg/kg/day with initial improvement but did not continue this more than a few weeks due to miscommunications. At 39 months of age, he was started on levetiracetam at 10 mg/kg/day then uptitrated to 40 mg/kg/day, providing partial control of his episodes. Repeat genetic testing was requested at 51 months of age as an epilepsy panel through Invitae but resulted only with variants of uncertain significance in *ARX*, *CPA6*, *FASN*, and *ST3GAL3*.

Patient 1 was seen by primary care at 4 years 11 months with seizures reported under good control but with continuing difficulty with speech. He was prescribed atovaquone/proguanil for malaria prophylaxis for an upcoming trip to Africa (Kenya and Somalia). During the trip, Patient 1 developed an illness and then had rapid progression of neurologic symptoms and respiratory failure attributed to “meningitis”. Unfortunately, he passed away aged 5 years, 2 months. Due to his passing away while abroad, limited information is available about the exact medical course of his passing.

### Patient 2

Patient 2 (II-2 in **Figure 1A**), sister and second cousin to Patient 1, is a female of Somali Ancestry born in Kenya. There were no complications with pregnancy nor birth history, and she was born full-term. Early milestones were recalled as being on time. Her language development was reportedly significantly delayed; she spoke single words at 24-26 months and was not able to produce two-word phrases until 42 months. Chart records begin at 34 months of age, 2 months after she arrived in the United States. At that time, she had a history of episodic emesis once to twice a month since a year prior. She was also noted to have granuloma annulare on her foot that eventually resolved. At 39 months, the family reported a 2-year history of shaking when she was sick. Urinary frequency and occasional incontinence with negative urinalysis and cultures began at 39 months. At 47 months of age, a neuropsychology evaluation diagnosed Mixed Receptive-Expressive Language Disorder. A murmur noted on a routine exam prompted an echocardiogram at 4 years old. However, the echocardiogram was normal and the murmur was not heard on later exams.

At 5 years old, difficulties with aggressive behaviors, severe enough to require withdrawal from school, prompted re-referral for developmental services. In conjunction with her brother's neurology visit at 39 months, she was evaluated at 6 years old by neurology for generalized seizures and 15-30 minute episodes of stiffening and weakness on either side of the body without loss of awareness. The seizures were twice per year and the stiffening episodes twice per month. Before age 3, seizures only occurred with fevers, but, afterwards, seizures began to occur without illness. Stiffening episodes were also noted to have started years prior to the seizures. She was started on levetiracetam 10 mg/kg/day and then uptitrated later to 50 mg/kg/day with good response of the stiffening episodes. An EEG at 7 years of age was abnormal due to presence of mild slowing of background interpreted as related to drowsiness but without clear epileptiform activity. Brain MRI at 8 years old was normal. At 8 years old, she was noted to have difficulty with slow transit constipation and started on polyethylene glycol.

Later that year and along with her brother, she was seen for a well check prior to a family trip to Africa (Kenya and Somalia). Atovaquone-proguanil malaria prophylaxis was prescribed to her as well. She returned 3 months later, presenting to the emergency room with 2-3 weeks of lower extremity weakness, urinary incontinence, and weight loss which all started in Somalia after 2 days of loose, watery, non-bloody stool. At the time of presentation, she was afebrile and the diarrhea had resolved. However, she was so weak that her parents had her in a diaper and using a wheelchair, no longer able to ambulate. She had pain in lower legs. Her speech was noted to be impaired and had decreased over the previous 3 weeks. Her deep tendon reflexes were present yet diminished. She was intubated for procedures, including a lumbar puncture and imaging, but was unable to be extubated due to diaphragmatic weakness. Her family described her illness course as being very similar to the process that had resulted in her brother's untimely passing. Patient 2's infectious workup was negative, with no indication of meningitis. MRI was interpreted as nonspecific and showed only a single focus of T2 hyperintensity within the periventricular white matter and no findings on spinal MRI. Lumbar puncture was acellular with normal glucose and protein. EEG was normal initially but subsequently showed subclinical seizures. Intravenous Immunoglobulin was given for possible Guillain Barré Syndrome even with present albeit diminished reflexes, and she also later underwent plasmapheresis without improvement. Nerve conduction study and electromyography 3 days into her admission showed electrodiagnostic evidence for likely an acquired and acute motor predominant peripheral polyneuropathy with possibly primary axonal involvement. Specifically, it showed markedly reduced compound motor action potential amplitudes in essentially all motor nerves (median, ulnar, tibial, peroneal) with mild and patchy involvement of sensory nerves. Fibrillation potentials were interpreted as being more likely neuropathic than myopathic due to slowing and sensory involvement. The study and clinical history were thought consistent with acute motor axonal neuropathy or acute motor and sensory axonal neuropathy.

Rapid quad exome analysis of genome sequencing was sent, including patient 2, mother, father and a residual DNA sample from her brother's (Patient 1) prior testing. This testing resulted with *ADPRS*:NM_017825.2; c.545A>G (p.His182Arg), homozygous and *ST3GAL3*:NM_006279.3; c.685G>A (p.Ala229Thr), homozygous. Both variants were heterozygous in each parent, homozygous in Patient 1, and assessed by the diagnostic lab as being of uncertain significance. On both a phenotypic basis and given the data available at the time, including *in vitro* characterization of a variant at the same amino acid residue, the *ADPRS* homozygous variant was felt to be the more likely explanation for the phenotype of both patients.

Due to persisting diaphragmatic weakness, Patient 2 underwent tracheostomy to allow long-term mechanical ventilation. Later a gastrostomy tube was also placed to facilitate feeding. She had a prolonged inpatient admission of 8 months due to inability to secure services in a home setting to support her ventilatory status. She developed neurogenic bowel and chronic pseudo-obstruction, requiring an aggressive bowel regimen. Influenza vaccine and 2 doses of COVID-19 vaccine were received without clinical exacerbation. She continued to have neuropathic pain symptoms requiring pregabalin, amitriptyline, gabapentin, and clonidine. Clobazam was added to levetiracetam for additional seizure control. Fosphenytoin was briefly used as well. Levetiracetam was later transitioned to lamotrigine. She was prescribed ramelteon for difficulty sleeping. During the admission, she had several bouts of tracheitis and urinary tract infection. She was ultimately diagnosed with a neurogenic bladder. At 9 years old, she had blurry vision and saw ophthalmology who noted astigmatism and prescribed glasses. Her hearing was checked by audiology and found to be normal.

She had multiple readmissions due to infections or complications of tracheostomy and ventilation. Neurological status seems to regress with each infection. She has continued to have progression of weakness and of verbal skills. She lost reflexes in both upper and lower extremities. During an admission at 10 years old, she developed a dyskinesia affecting her face and upper arms, with legs completely denervated by this point. This dyskinesia was attributed to concurrent use of levofloxacin and nitrofurantoin in the setting of her vulnerable nervous system. Over time she developed temperature dysregulation and dysautonomia. Her neuropathy progressed to quadriparesis and generalized hypotonia. She developed knee contractures. and neuromuscular scoliosis. With immobility, she developed deep vein thrombosis, osteopenia, and subsequent bone fractures. She continued to have severe insomnia. Seizure control became more difficult, although much of her seizure activity remained subclinical. Electroencephalography showed generalized abnormalities implying diffuse bilateral cortical dysfunction. Seizure control adjustments continued with discontinuation of clobazam and initiation of phenobarbital and transition back to levetiracetam from lamotrigine. At 11 years old she developed total cataracts of both eyes.
